# Deep learning and cross-species analysis identify the influences of sex and cannabinoid signaling on cerebellar vermis morphology

**DOI:** 10.1101/2021.07.03.450642

**Authors:** Kylie Black, Om Pandey, Brent McPherson, John Hainline, Shawyon Baygani, Brynna Webb, Tristen Mier, Amanda Essex, Ken Mackie, Franco Pestilli, Anna Kalinovsky

## Abstract

The cerebellum is the only folded cortical structure present in both rodents and humans, allowing mechanistic studies of gyrification across species. In mice, the extent of anterior vermis folding and the shapes of white matter exhibit sex-dependent differences. We trained a deep convolutional neural network to predict biological sex from midline cerebellar MRI images, showing that human cerebellar midvermis contains sex-dependent anatomical features. Our comparative cross-species analysis shows that in both species, sex and anterior vermis folding patterns are not enough to predict an individual’s behavior because the variability in performance between individuals is far greater than the differences between conditions. Finally, utilizing constitutive cannabinoid receptor 1 (CB1) knockout (KO) mice, we identify developmental cannabinoid signaling as a novel molecular mechanism limiting secondary fissure formation.

**Teaser:** Males and females display differences in the architecture of cerebellar folding shaped by cannabinoid signaling during development.

## Introduction

From ancient times, humans have been fascinated by the intricate patterns of brain folding and how it may correlate with differences in individual behaviors: ancient Egyptian papyrus transcribed in the 7^th^ century BC contains a description of the folded brain appearance “like ripples that happen in copper through smelting” (1). In colloquial Russian, “missing a few brain folds” implies that the person is not very smart. In fact, variability in cortical folding accounts to a large extent for the individual variability in brain anatomy (2). However, genetic and signaling mechanisms regulating the development of folding patterns are not well understood. Even though the anatomical boundaries of lobules do not always correspond with the boundaries of functional cerebellar subregions (3), they create physical constraints on neuron migration and the development of neural microcircuits (4). Variability in cerebellar folding has been linked to differences in behavioral outputs in human neurodevelopmental disorders (5) and in the variability of behavioral complexity between species (6).

The cerebellum is a tightly folded cortical structure, its flattened area equaling about 80% of the area of the human cerebral cortex, yet it contains a greater number of neurons (7) and is implicated in a wide range of cognitive, emotional, and motor behaviors (8, 9). It exhibits intricate anatomical and functional compartmentalization, including rostrocaudal folding of the vermis into ten primary lobules, which are conserved across vertebrates, and secondary lobules, which are more variable between species and across individuals within species (10). The cerebellum is not considered sexually dimorphic, albeit alterations in cerebellar structure and function in humans have been associated with neuropsychiatric disorders that exhibit higher prevalence or earlier onset in males, including autism spectrum and schizophrenia (11).

Here, we report that the prevalence of secondary fissures in the anterior cerebellar vermis differs between males and females, identifying the cerebellum as a brain region exhibiting sex-dependent morphological differences in both mice and humans. We show a correlation between differences in the foliation patterns and the shapes of the white matter (WM) tracts. However, we did not find either sex or the differences in cerebellar vermis foliation sufficient to predict the performance of gross or fine motor tasks. The differences in motor tasks performance between individuals are much greater than the mean differences between sexes, underscoring that neither sex nor anatomical differences are enough to predict an individual’s behavior. The expression of CB1 cannabinoid receptors in the anterior vermis of mice is high at birth and decreases during the first postnatal week (12) when secondary fissures are initiated. Here, we show that CB1 expression adjusts the timing of formation and the abundance of secondary fissures in mice, highlighting a previously unrecognized role of cannabinoid signaling as a molecular mechanism regulating cortical folding.

## Results

### The Predominant Pattern of Vermis Foliation is Sex-Dependent in CD1 Mice

In his foundational comparative analysis of cerebellar foliation, Olof Larsell identified ten primary vermal lobules (I-X) conserved in most vertebrates (10). Using Larsell’s classical anatomical criteria, lobules II through X can be consistently recognized in all CD1 mice (Fig. 1 A-A”). Per Larsell’s nomenclature, lobe I is a half-lobe nested on the dorsal surface of the anterior medullary velum and is characterized by its white matter resting directly on the velum. Guided by this definition, lobe I (Fig. 1 A” – green arrowhead marks the fissure separating lobes I and II) is extremely rare in CD1 mice (1/100 in our colony – in contrast to the C57B6 inbred strain that was not used in our study). In the midvermis of CD1 mice, we identified three hotspots with high variability between individuals: (1) fissure *precentral-a* (*prc-a*) separates lobules IIa and IIb (Fig. 1 A’), while in other animals, *prc-a* is absent leaving lobule II undivided (Fig. 1 A, A”); (2) *intra- culminate* (*ic*) fissure, separating lobules IV and V, is present in some (Fig. A, A’) and absent in other (Fig. A”) animals; (3) in some animals a *superiordeclive* (*s-de*) fissure is present dividing lobules VIa and VIb (Fig. 1 A, A”), but not in (Fig. 1 A’). Sites of these secondary fissures are marked by green-lettered abbreviations and with asterisks where the fissures are present (Fig. 1 A-A”). Anatomical nomenclature was adopted from (10, 13).

**Fig. 1.**
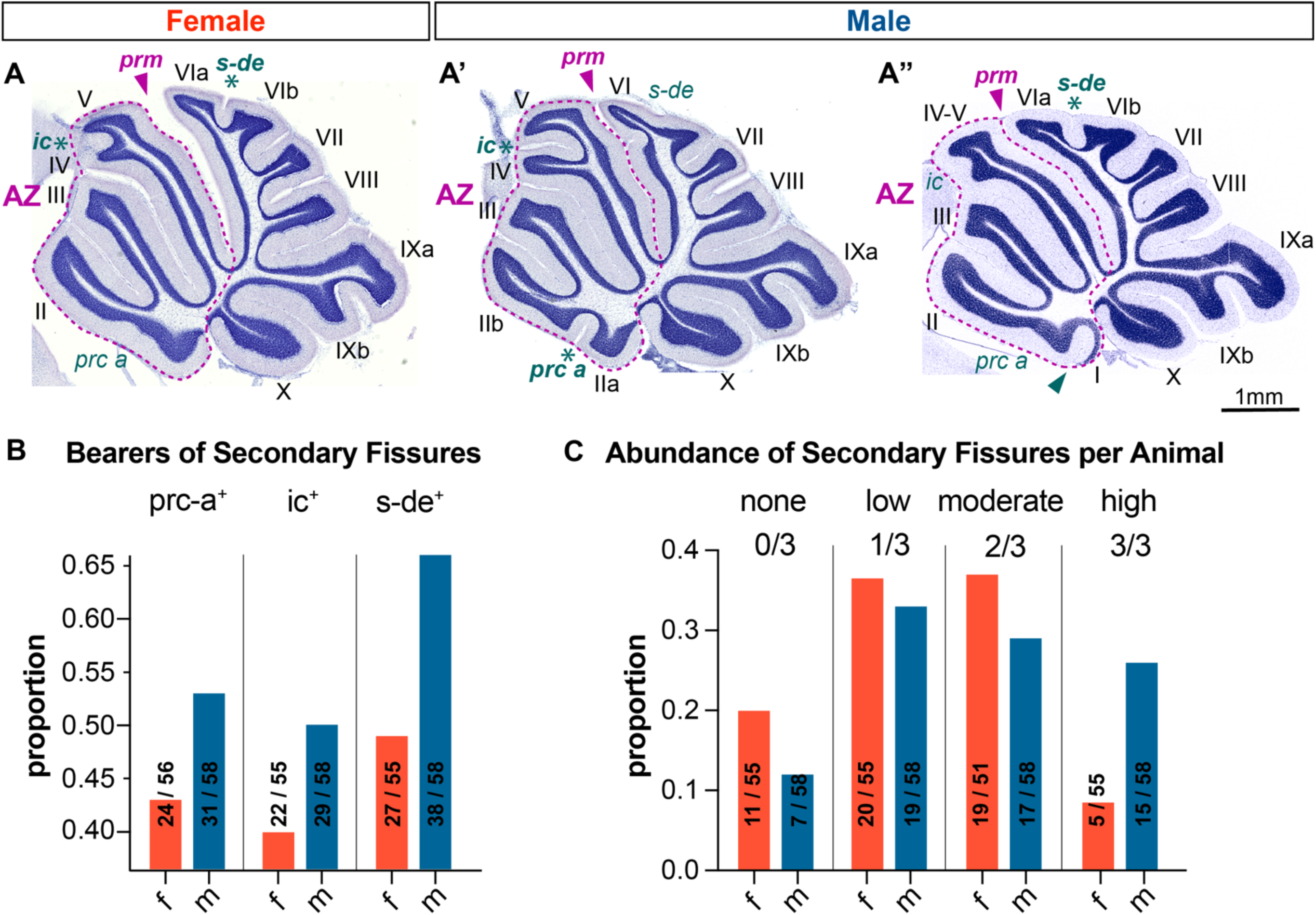
Secondary fissures in the cerebellar vermis are more prevalent in the male CD1 mice. Representative female (**A**) and male (**A’**) midsagittal cerebellar sections (rostral = left, dorsal = top). Fissures separate the vermis into lobules (designated by Roman numerals). Locations of secondary fissures are shown by green-lettered abbreviations and by an asterisk when the fissure is present. Purple arrowheads mark the primary fissure (*prm*), a landmark designating the boundary of the anterior zone (AZ) – outlined in purple. (**A”**) Shows a rare case where lobe I, a half-lobe nested on the dorsal surface of the anterior medullary velum, can be seen. A fissure separating lobe I from lobe II is marked by a green arrowhead. (**B**) The proportion of animals expressing each one of the secondary fissures (*prc-a*, *ic*, and *s-de*) is higher in males, and a higher proportion of males express all three secondary fissures (**C**). 56 female and 58 male WT CD1 mice were analyzed at two months of age.

Since secondary fissures are variable between individuals, we asked what factors governing cerebellar development may increase the probability of secondary fissure formation. A ubiquitous genetic mechanism contributing to variability between individuals within vertebrate species is genetic sex. Thus, we examined the relationship between sex and the prevalence of secondary fissures. 56 female and 58 male WT CD1 mice were analyzed at two months of age. Our results show that in two-moth-old young-adult mice, more males than females exhibit any of the three most common secondary fissures: *prc-a^+^* – 53% of males vs. 43% of females, *ic^+^* – 50% of males vs. 40% of females, *s-de^+^* – 66% of males vs. 49% of females (Fig. 1 B). Males are more likely to express multiple secondary fissures (males are 3 times more likely than females to express all three secondary fissures, while females are twice more likely than males to express no secondary fissures at all, Fig. 1 C). Thus, the cerebellar vermis exhibits sex-dependent foliation differences in CD1 mice – males are characterized by a higher probability of secondary fissure expression and a higher total number of secondary fissures per animal.

### Human Sex is Well Predicted Using Cerebellar Anatomical Images

The high degree of variability in midvermal foliation in humans was emphasized by Olof Larsell based on postmortem examination of human brain tissue in (Fig. 2 A, from “Human Cerebellum” figure 54, p.41; (10)). It is also evident in MRI scans from the Human Connectome Project (HCP, https://www.humanconnectome.org/ – Fig. 2 B). For that reason, and because the imaging resolution of MRI scanning is lower compared to classical histological staining and microscopy, it is impractical to conduct the same type of manual anatomical classification that we employed for the mouse cerebella. Thus, to investigate whether, in humans, the foliation pattern of the cerebellar vermis is influenced by sex, we chose to use an unbiased approach to probe for anatomical differences.

**Figure 2.**
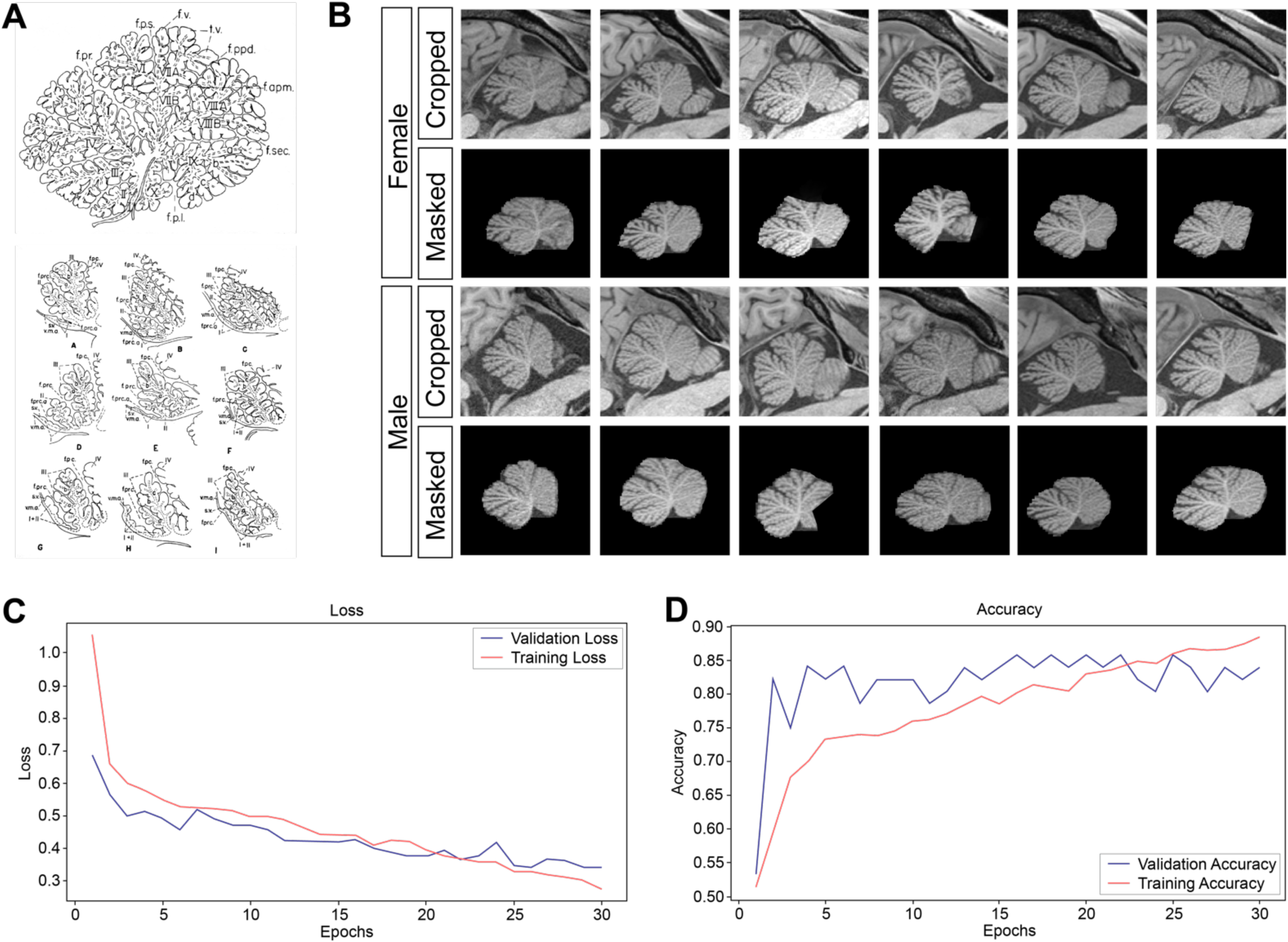
Human sex is well predicted using cerebellar anatomical images. **(A)** Traced midsagittal human cerebellar vermis with lobules and fissures marked (lobules I-X, rostral = left, dorsal = top; the primary fissure separating anterior zone is abbreviated **f. pr**.) from Olof Larsell’s ***The Comparative Anatomy and Histology of the Cerebellum, Human Cerebellum,*** figures 55 B, p.45 and 54, p.41 (Larsell, 1967) illustrating the high degree of variability in anterior vermis folding. **(B)** Representative examples of midline sagittal cerebellar images from HCP. Images are shown in the same orientation as in (A). Top rows: images were selected by registering labels from the SUIT atlas to the subjects’ space and passed through an intensity normalization process that scaled the images for training (J. Diedrichsen et al., 2009; J. Diedrichsen, 2006; J. Diedrichsen et al., 2011). Bottom rows: in order to limit analysis to the midvermal cerebellum images were masked, all pixels outside the label set were zeroed, and images were cropped centered on the cerebellum, resulting in image sizes of 420, 280, 1 (height, width, channels, respectively). **(C)** A deep convolutional neural network (CNN) was trained to predict biological sex from cerebellar midvermis anatomy. Loss of prediction accuracy over time during training was used to evaluate model performance. **(D)** The model was run for 30 epochs with a batch size of 32 for training. The prediction accuracy was used to score model performance. Maximum accuracy achieved is 85.71% on validation and 88.37% on training, showing that the model has not overfit and can predict biological sex from anatomical cerebellar midvermis scans with high accuracy. 1114 sagittal cerebellar MRI scans from the Human Connectome Project (HCP, https://www.humanconnectome.org/) were used.

We reasoned that if there are mesoscale anatomical differences in cerebellar vermis morphology between human males and females, it should be possible to train a convolutional neural network (CNN) to determine biological sex from midvermal cerebellar scans. To test this hypothesis, we trained CNN to differentiate two categories: male versus female, based on isolated cerebellar midsagittal vermis MRI scans from the Human Connectome Project (HCP – 1114 images in total, split into 85%, 3%, and 12% for training, validation, and testing, respectively). Labels from the SUIT atlas were moved to the subject’s native space after alignment to identify the midline sagittal plane (14–16). Midline sagittal images were cropped and masked in order to limit the analysis to the midvermal cerebellum (Fig. 2 B). All pixels outside the midline cerebellar area were zeroed so no confounding information from surrounding tissue would contribute to the prediction, reducing image sizes from 1167, 875, 3 pixels (image height, image width, and the number of channels, respectively) to 420, 280, 1, by cropping the images around the cerebellum and converting from 3 (RGB) channels to single (grayscale) channel. We conducted batch training, with a batch size of 32 on multiple CNN models to find the optimal model size, using an Adam optimizer to predict 0/1 for male/female respectively and to experiment with the learning rate and threshold of validation. Data splits were independent of one another, and the model was validated every 5 epochs. Training versus Validation loss over epochs was plotted to track how the confidence of our model is changing during training (Fig. 2 C).

Images were passed through multiple layers of convolutional filters. The filters initially describe the local patterns found in the image, but as the layers later in the model become smaller, they represent more global properties of the image. The decision layer used the previous layers’ combined weights to make the desired category assignment. The network was trained for 30 epochs. Since there is not much difference in training data and the maximum accuracy achieved is 85.71% on validation and 88.37% on training, we conclude that the model predicts biological sex with high accuracy and is not overfit (Fig. 2 D). Thus, in humans, as well as in mice, cerebellar vermis exhibits sex-dependent morphological differences.

### In CD1 Mice, Differences in Vermal Foliation are reflected in the Shapes of Interlobular White Matter Tracts but are Independent of Vermal Size

Increased foliation and larger cortical size have been positively correlated in prior studies (17–19). In order to elucidate whether overall increased foliation in males may be linked to increased cerebellar vermis size, we measured the size of the midvermal area and found it to be slightly larger in males (Fig. 3 A – 11 mm^2^ mean in females and 12 mm^2^ mean in males, P-value=0.02). However, this difference of means between sexes is many folds smaller than the distribution of values within each sex (from less than 8 to about 16 mm^2^) and is erased when the midvermal area is normalized to animal weight (Fig. 3 B). We found no differences in midvermal sizes in animals with *prc-a* (prc-a^+^) or without (prc-a^-^) in both sexes (Fig. 3 A). Since the majority (two out of the three variable secondary fissure hotspots) of foliation differences localize to the anterior zone (AZ, lobules II-V, rostral from the primary (*prm*) fissure, purple dotted outline in Fig. 1 A-A”), we also assessed the differences in the anterior zone area and the ratio of the anterior zone to total midvermis area between animals with or without *prc-a* but found none (Fig. 3 C, D). In conclusion, the prevalence of secondary fissures does not correlate with increases in total or anterior midvermal area, suggesting that the increased incidence of secondary fissures is independent of the increase in cortical size.

**Figure 3.**
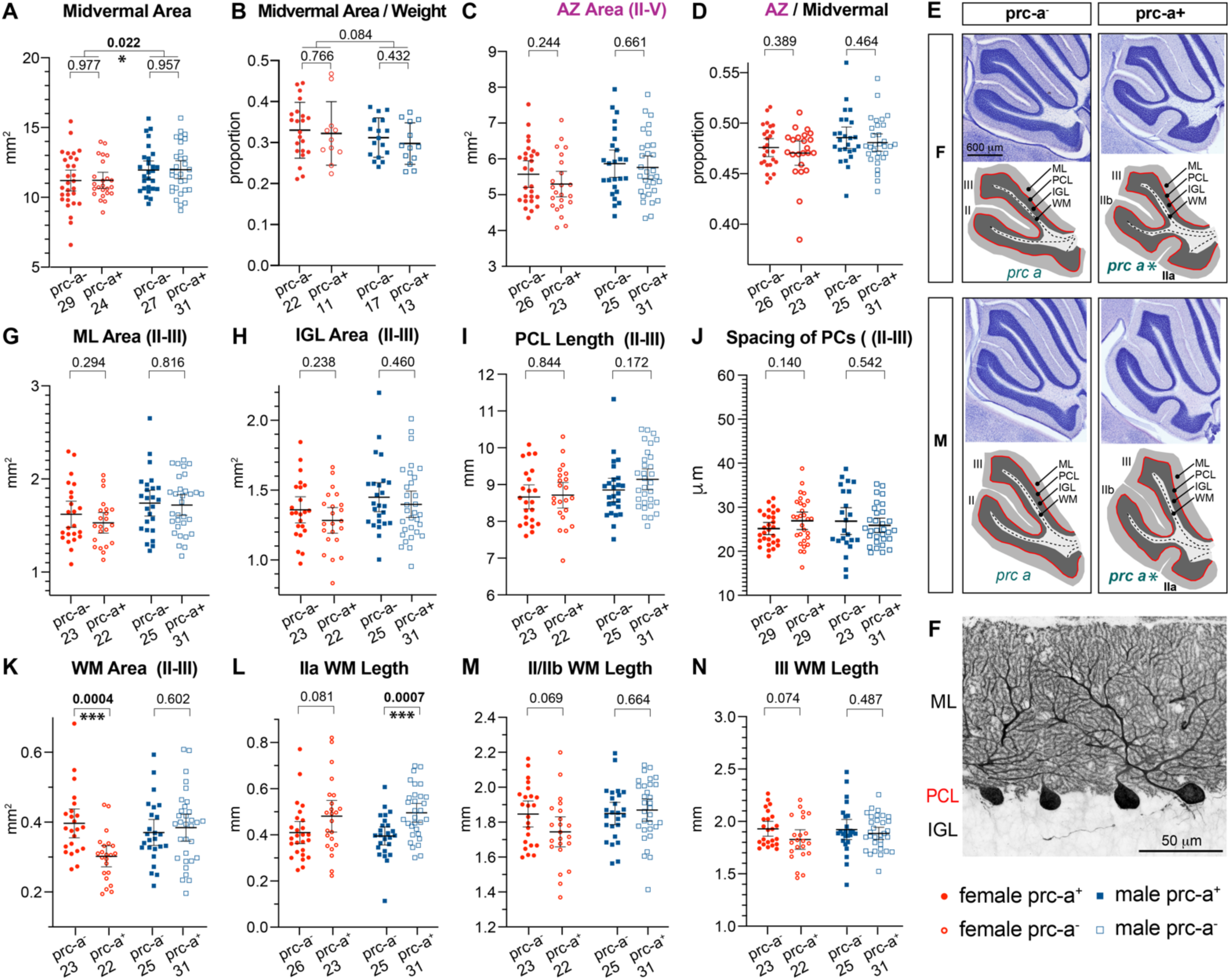
Foliation pattern affects white matter morphology but is independent of vermal size. **(A)** Total midvermal areas range from about 5 mm^2^ to over 15 mm^2^ in two- month-old CD1 mice of both sexes. For either sex, there are no differences in total midvermal size between animals expressing the secondary fissure *prc-a* (prc-a^+^) and those that do not express *prc-a* (prca^-^). The average size of midvermal area is about 1 mm^2^ larger in males (P=0.02), but the difference between sexes in total midvermal area sizes is no longer apparent when **(B)** midvermal area is normalized to animal weight. **(C)** Anterior zone area and **(D)** ratio of anterior zone out of total midvermal area do not differ between prc-a^+^ and prc-a^-^ animals. **(E)** Representative midsagittal sections showing lobules II-III in the anterior zone (AZ) of cerebellar vermis. Diagrams highlight cerebellar layers: molecular layer (ML), Purkinje cell layer (PCL), inner granule cell layer (IGL), and white matter (WM). Punctate lines trace the course through the middle of WM tracts to the folia apexes that was used to measure the lengths of WM tracts. In prc-a^-^ animals, where no distinct lobe IIa is present, the length of IIa WM tracts was estimated by tracing to the apex of the WM that bulges out in the corresponding place. Folia are designated by Roman numerals, sites of the secondary fissure *prc-a* by green lettered abbreviations, and asterisks indicate when the fissure is present. **(F)** Zoomed-in view of PCL on the ventral side of lobe II. Calb immunolabeling was used to visualize PCs for quantification of PC density per PCL length. **(G-J)** From midsagittal sections lobes II-III, quantification of areas of cerebellar molecular (ML) and inner granule cell (IGL) layers, lengths of Purkinje cell layer (PCL) and spacing of Purkinje cells (quantified from ventral side of lobe II) show no differences between sexes. **(K-N)** Quantification of areas and lengths of WM fascicles projecting into individual folia in lobules II-III. Abbreviations: AZ = anterior zone, IGL = inner granule cell layer, ML = molecular layer, PCL = Purkinje cell layer, WM = white matter, *prc-a* = anterior precentral fissure.

The cerebellar cortex is composed of morphologically, cellularly, and functionally distinct layers: the outermost molecular layer (ML), underlied by the Purkinje cell layer (PCL), with the inner granule cell layer (IGL) located in the innermost portion of the cerebellar gray matter, resting on top of the white matter (WM). Overall shapes of lobules II-III are distinctly different in prc-a^+^ mice (resembling a trefoil leaf) versus prc-a^-^ mice (two-prong fork shape) (Fig. 3 E). However, it is not clear whether these differences in the general shape reflect differences in the size and composition of the gray and white matter. To evaluate what structural elements of the cerebellar cortex are affected by the expression of secondary fissures, we quantified areas of the layers in lobules II-III from midsagittal sections (Fig. 3 E – IGL is shaded dark gray and ML light gray in the masked regions of interest corresponding to the representative Nissl-stained sections), the length of the PCL (Fig. 3E – red line). The average spacing between Purkinje cells (PCs) within the PCL was quantified from the higher magnification images taken from the ventral side of lobe II (Fig. 3 F). We found no significant differences between prc-a^-^ and prc-a^+^ in the mean ML area (Fig. 3 G), the mean IGL area (Fig. 3 H), PCL length (Fig. 3 I), or spacing of PCs (Fig. 3 J). Thus, the differences in the overall shapes of lobules II-III do not correspond to differences in the sizes of the layers composing cerebellar gray matter.

We measured the total areas and lengths of WM fascicles projecting into individual folia (IIa in prc-a^+^ or the apex of the corresponding WM bulge in prc-a^-^; IIb; and III (Fig. 3 E – WM area is masked with the light gray, the trajectories tracing the middle of the fascicle that have been used to evaluate fascicle lengths are shown as punctate lines). In prc-a^+^ females, the overall area of WM in lobules II-III is smaller (Fig. 3 K, P-value=0.0004), matched with a trend for longer tract IIa (Fig. 3 L, p=0.081) and shorter IIb (Fig. 3 M, p=0.069), corresponding with the generally stockier trefoil shape of lobules II-III and more branched WM. In prc-a^+^ males, the overall shape of WM is also stockier, and IIa is significantly longer (Fig. 3 L, P-value=0.0007), with no significant differences in II-III WM area or IIb length. In conclusion, increased vermal folding is associated with a stockier overall shape of lobules II-III, corresponding to structural changes in cerebellar WM tracts that are more branched but slimmer. Increased vermal folding does not affect the areas of neuronal layers or the numbers or spacing of PCs.

### Vermis Foliation and Sex Have no Decisive Impacts on Behavior

The anterior cerebellar vermis receives projections from the motor, somatosensory, and vestibular afferents and is activated during tasks that involve posture, locomotion, and forelimb dexterity (20, 21). We wondered whether mesoscale anatomical differences in the anterior cerebellar vermis – distinct folding patterns and WM shapes – are associated with detectable differences in behaviors. Since humans exhibit sex-dependent anatomical differences in the anterior cerebellar vermis, we asked whether there is sexual dimorphism in the performance of anterior cerebellum-associated behaviors in humans. Behavioral data from HCP includes gait analysis (time to walk the length of a corridor) and forelimb dexterity task (placing pegs in holes) (Fig. 4 A, B – 202 females and 303 males). The performance in the gait task is comparable between sexes (Fig. 4 B). In the peg sorting task, males are slightly faster than females (Fig. 4 A, the mean for females is 110.7 seconds, and for males 114.4 seconds; differences between means ± SEM = 3.713 ± 0.9414, P-value < 0.0001, as determined by an unpaired T-test). Importantly, this analysis also highlights the wide variability in performance between individuals, which is many folds greater than the slight differences in mean performance between sexes.

**Figure 4.**
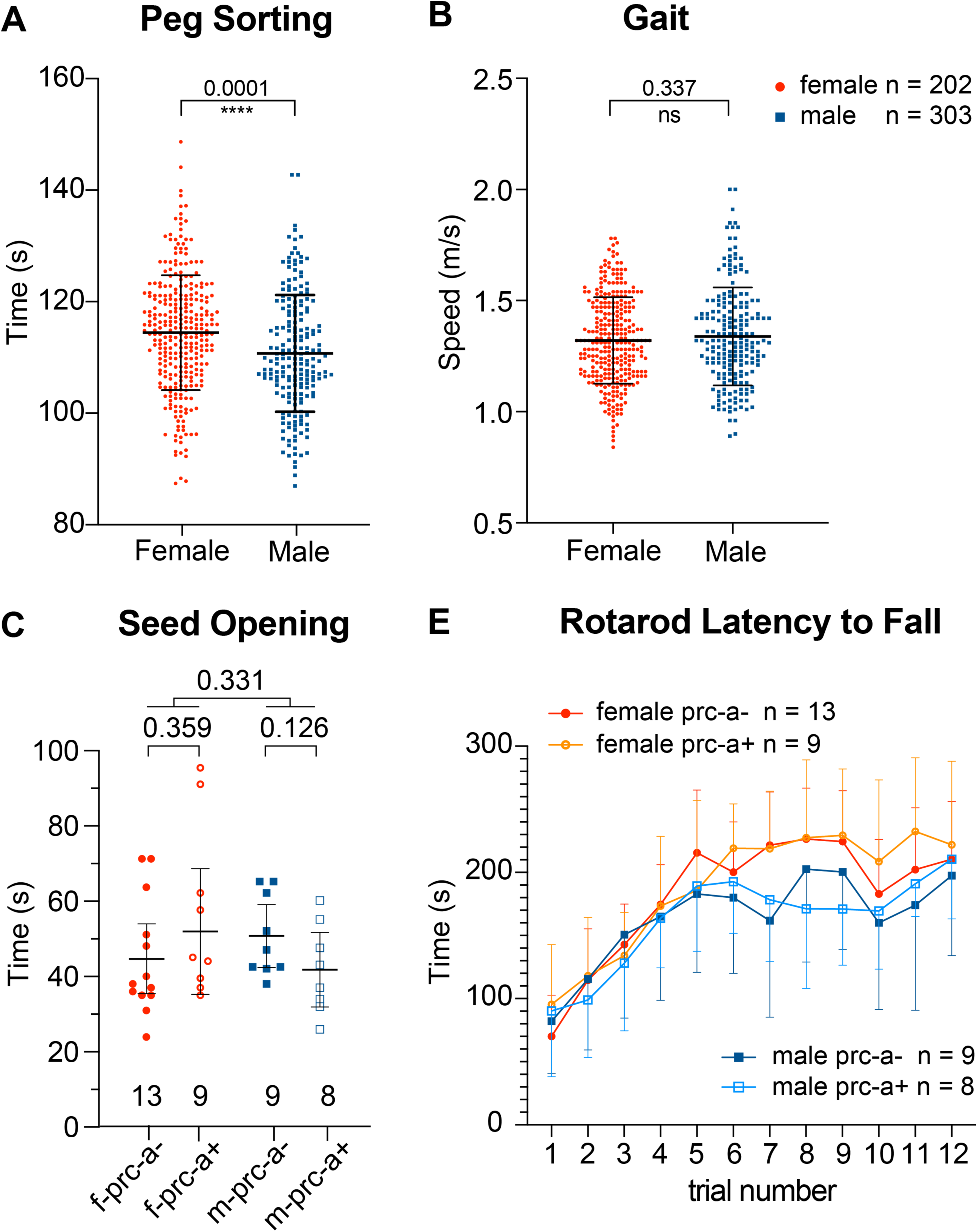
Neither Anatomical Differences in Anterior Vermis Foliation nor Biological Sex Exhibit a Definitive Influence on Behavior. (A) Human subjects performing peg sorting forelimb dexterity task (data from HCP). The Y-axis shows the time (in seconds) required to accurately place and remove 9 plastic pegs into a plastic pegboard with a dominant hand. **(B)** Gait analysis of human subjects (data from HCP). The Y-axis shows the average speed of locomotion (meters per second) assessed after the subjects walk the length of a corridor. **(C)** Forelimb dexterity in mice was assessed by the sunflower seed opening task. There are no statistically significant differences between sexes or depending on the presence of prc-a. The presence of prc-a is associated with opposing trends in females vs. males: slower mean time to open seeds in females and faster in males. Columns show means +SD. An unpaired student T-test was used to assess P-values. **(C)** The expression of prc-a does not affect rotarod performance.

We also assessed sensorimotor coordination in a rotarod test and forelimb dexterity in a sunflower seed opening task in CD1 mice and detected no significant differences between sexes (Fig. 4 C, D). Since the presence of secondary fissures is more common in males, could the influence of sex on seed opening be directly linked to a higher degree of anterior vermis foliation? Looking at the correlation between seed opening times and the presence of *prc-a* (Fig. 4 C), we found opposing trends between sexes but no statistically significant differences. In conclusion, both biological sex and the extent of anterior vermis foliation do not exhibit a detectable influence on behavioral performance.

### The Timing of Secondary Fissures Initiation Contributes to the Sex Differences in their Prevalence

Primary fissures, which are highly conserved between individuals, are initiated early in development, deepening and becoming more pronounced while the cerebellar cortex undergoes its developmental expansion. In contrast, secondary fissures are initiated later, are less pronounced, and vary in prevalence between species and individuals (22–24). Characterization of the developmental timecourse of secondary fissure formation in WT CD1 mice shows that *prc-a*, *ic*, and *s-de* are initiated after postnatal day 3 (P3) (Fig. 5 A, B, C-D). In males, a consistently higher percentage of secondary fissures are detected at earlier time points than in females: ***prc-a*** – at P5 10% of males and none of females exhibit *prc-a*, and the proportion of prc-a^+^ animals increases with age in both sexes, reaching ∼53% in males and ∼43% in females at two months of age (2mo) (Fig. 5 C); ***ic*** – at P5 ∼30% of males and ∼20% of females express *ic*, reaching ∼50% in males and ∼40% in females by P10-2mo (Fig. 4 D); ***s-de*** – at P5 ∼40% of males and ∼25% of females express *ic*, and by P10 the maximum proportion of s-de^+^ (∼75%) is reached for both sexes (Fig. 5 E). Thus, the higher prevalence of secondary fissures in males may be a consequence of an earlier and longer developmental window for secondary fissure initiation.

**Figure 5.**
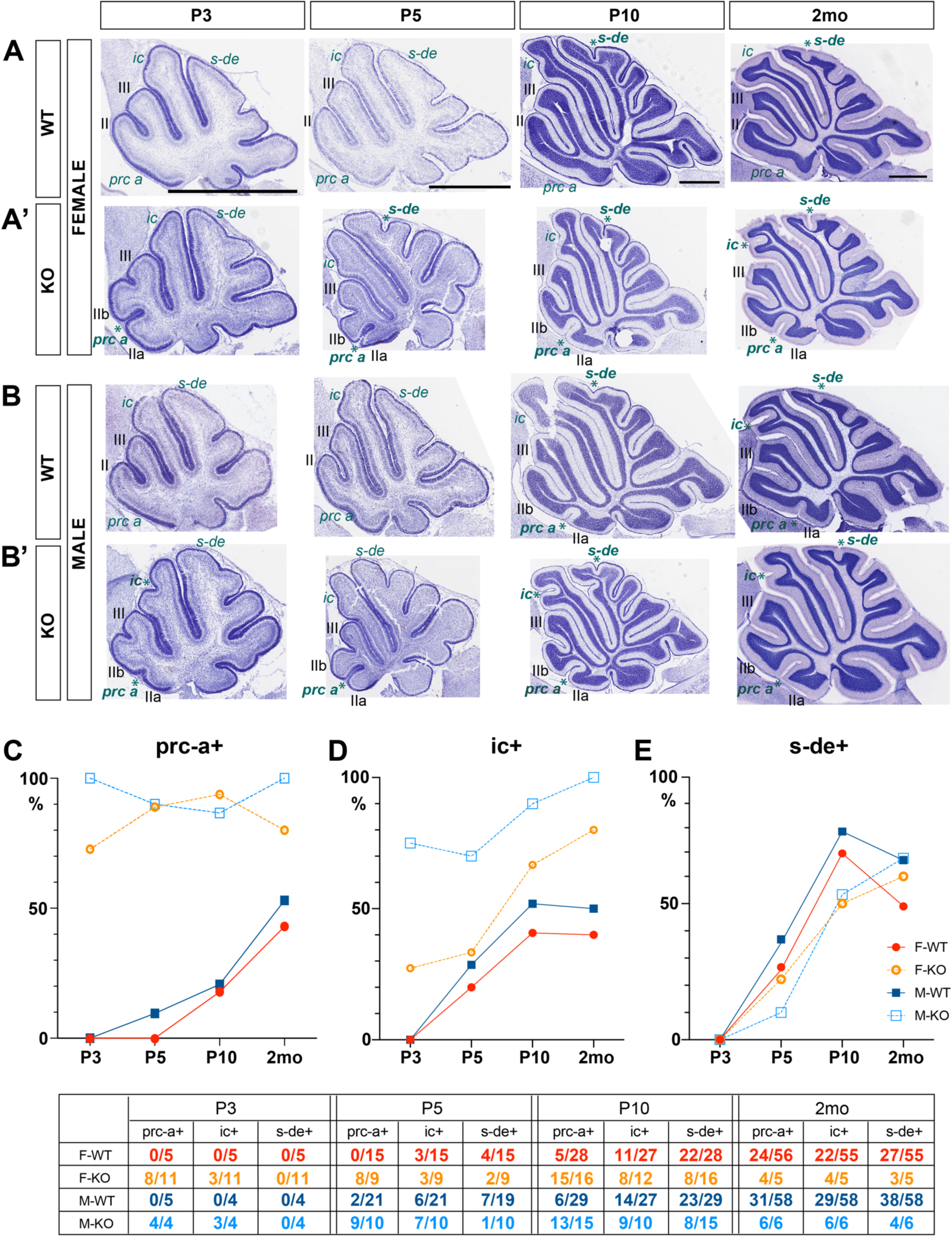
Cannabinoid Signaling Regulates Development of Secondary Fissures. Representative midvermal images from CD1 mice showing developmental timecourse of secondary fissure formation in **(A) f**emale wildtypes, **(B)** male wildtypes, **(A’) f**emale knockouts, **(B’)** male knockouts. **(C)** Quantification of the proportion of animals expressing ***prc-a*** at the different developmental timepoints. **(D)** Quantification of the proportion of animals expressing ***ic*** at the different developmental timepoints. **(E)** Quantification of the proportion of animals expressing ***s-de*** at the different developmental timepoints. Abbreviations: *ic* = intraculminate fissure, *pcen* = precentral, *prc-a* = anterior precentral fissure, *prm* = primary fissure, *s-de* = superior declive fissure, F = female, M = male, P = postnatal day.

### Cannabinoid Signaling Regulates Development of Secondary Fissures

In our work with constitutive CB1 cannabinoid receptor knockouts (CB1-KOs), we noticed an increased incidence of secondary fissures (Fig. 5 A’, B’). Strikingly, the majority of CB1-KOs of both sexes exhibit anterior zone secondary fissures *prc-a* and *ic* (Fig. 5 A’, B’).

Conversely, there is no difference between WTs and CB1-KOs in the prevalence of *s-de*, located in the central zone (Fig. 5 A’, B’). Furthermore, *prc-a* and *ic* are initiated prematurely in CB1 KOs: by P3 ∼100% in males and ∼73% of females are prc-a^+^, and ∼75% of males and ∼27% of females are ic^+^ (Fig. 5 C, D). These observations suggest that cannabinoid signaling through CB1 limits the formation of secondary fissures by delaying the developmental window during which secondary fissures are initiated. Consequently, increased foliation due to the common occurrence of secondary fissures in the anterior zone is a distinguishing morphological feature of CB1 knockouts.

## Discussion

In sexually dimorphic brain regions, such as the medial preoptic area of the hypothalamus, sex is an important biological variable affecting multiple aspects of nervous system development – from regulation of programmed cell death to synaptic pruning – therefore affecting sizes of brain regions and neuronal populations, neuronal and neuroendocrine activity, and synaptic connectivity (reviewed in (25)). However, the developmental effects of sex on sculpting cerebellar anatomy have not been previously described. Therefore, we were surprised by our initial observation in mice that the prevalence of secondary fissures, and thus the predominant folding patterns in the anterior cerebellar vermis, differ between sexes. Our results show that males exhibit extra-numeral fissures compared to females in WT mice. To address whether humans also exhibit sex-dependent variability in cerebellar vermis anatomy, we used a large open-access MRI database containing brain images from more than 1000 healthy subjects (HCP). Limiting the analysis to the cerebellum, we trained a 2D convolutional neural network on midsagittal anatomical MRI scans and achieved >85% accuracy in predicting biological sex in humans. Our cross-species analysis demonstrates that in both mice and humans, biological sex influences folding patterns of the cerebellar vermis, expanding the cohort of brain regions known to harbor sex-dependent anatomical differences.

### Morphometric differences in the white rather than gray matter underlie the different foliation patterns

The major cerebellar neuronal cell types are arranged in distinct layers. We performed morphometric analysis of the WM and the cortical gray matter layers in lobules II-III of the cerebellar vermis in mice. This analysis did not reveal any differences in the areas of any of the cortical layers in *prc-a^+^* versus *prc-a^-^*mice. Furthermore, the numbers and density of PCs, the principle cerebellar neurons that dictate the firing patterns of cerebellar outputs, are unaffected by the different foliation patterns, since neither the overall length of PCL, nor the spacing of PCs, differ in lobules II-III between *prc-a^+^*and *prc-a^-^* mice. We conclude that the differences in the overall shapes of the anterior cerebellar vermis most likely do not affect the numbers or density of neurons composing these layers. In contrast, our morphometric analysis identified sex and foliation-dependent differences in WM volumes in lobules II-III. Both sexes exhibit similar trends towards slimmer and longer tracts, with a statistically significant reduction in WM tract volume in females and statistically significantly longer tracts in males. Our study highlights the differences in WM volume as the most prominent morphometric distinction between prc-a^+^ and prc-a^-^ mice. This suggests a possibility that the mechanical forces regulating WM development could contribute to cortical folding. To result in slimmer WM tracts, are the axons in the developing cerebellar WM more fasciculated during development? Are there differences in the numbers of oligodendrocytes or the extent of myelination in the WM between prc-a^+^ and prc-a^-^ mice that can result in differences in WM volume?

### Examining the architectonic hypothesis – do the differences in the folding patterns correspond to distinct cytoarchitecture and behavior?

Advancements in the development of non-invasive brain imaging methodology and the computational tools for anatomical shapes analysis have renewed interest in the age-old fundamental neuroscience question of shape-function relationships and the hope to find non-invasive imaging biomarkers that can be utilized in the diagnosis and prognosis of neurological and psychiatric conditions. A logical assumption is that the sizes and shapes of brain regions create anatomical constraints to the number of cells and the patterns of neuronal connectivity. This hypothesis suggests a causal relationship between mesoscale anatomical differences in cortical foliation patterns, underlying cytological differences in distinct configurations of microcircuits, and individual variability in functional and behavioral outputs. There is high individual variability in both the cerebellar foliation patterns and cerebellar-influenced behaviors. We asked whether there is a correlation between the individual variability in the folding patterns and the behavioral outputs.

We observed no statistically significant differences between sexes in sunflower seed opening and rotarod tasks in mice. Furthermore, if the trend for males to slightly outperform females in seed opening (about 10 seconds shorter mean time to open a seed) corresponds with a higher proportion of males expressing *prc-a*, we expected prc-a^+^ males to outperform prc-a^-^ males in seed opening – this expectation is not supported by the data. Analysis of human gait and peg sorting data from HCP database shows a similar absence of decisive influence of biological sex on cerebellar-vermis associated behaviors – the mean differences between males and females are small, and even though unpaired T-test assigns significant P-value to the differences between males and females, the mean difference in the performance of peg sorting task is many fold smaller than the differences between individuals in each group. In summary, neither sex nor anatomical differences in the anterior cerebellar vermis provide predictive behavioral markers that we could detect.

### Endocannabinoid signaling regulates secondary fissure formation

Endocannabinoid signaling plays an important role in brain development (26), and early life exposure to phytocannabinoids is considered a risk factor for the development of schizophrenia, emotional and attentional deregulation, and substance use disorder (27). The most abundant neuronal cannabinoid receptor, CB1, is highly expressed in the developing axon tracts in the anterior vermis of newborn mice, but its expression is rapidly downregulated during the first postnatal week (12), during the developmental window of *prc-a* and *ic* initiation. Here, we show that in CB1 KO mice, the developmental window of *prc-a* and *ic* initiation shifts earlier, with both males and females expressing super-numerous secondary fissures. These results suggest that cannabinoid signaling regulates the timing of initiation and the abundance of secondary fissures, highlighting a previously unrecognized role of cannabinoid signaling as a molecular mechanism adjusting cortical folding.

The sex-dependent differences in anterior vermis foliation suggest that circulating steroid hormones or other effectors of genetic sex differences regulate CB1 expression during the perinatal period of secondary fissure initiation. Since secondary fissures are initiated earlier in males, it will be interesting to test in the future whether higher levels of circulating testosterone accelerate the downregulation of CB1 expression in the developing cerebellar axon tracts in males. Additional future questions include the effects of perinatal exposure to plant-derived cannabinoids on cerebellar foliation.

### How do the results of this work fit with the existing hypotheses of cortical folding?

Three distinct hypotheses have been proposed to explain the mechanisms regulating cortical folding. First is the fitting hypothesis stating that increased cortical folding correlates with developmental and evolutionary increases in cortical areas, and the folds develop due to mechanical crumbling, allowing the expanding surface to fit in a confined space limited by cranial bones (for example, (28)). Our results contradict this hypothesis since CB1 KOs exhibit both decreased cortical size (12) and increased foliation.

The second hypothesis suggests that differential expansion of the cortical layers leads to asymmetrical cortical growth, creating folia. Prior studies have focused on the cerebellar surface and the GC layer as the origin of the mechanical forces that drive cerebellar folding. Initiation of fissure development has been proposed to depend on the formation of developmentally transient structures: fissure anchoring points, generating physical constraints of cortical expansion by constraining GC migration through tight junctions between the endfeet of Bergman glia and the invaginated basilar membrane (22–24). Our results do not directly support this hypothesis since we did not identify differences in gray matter areas in the anterior vermis between prc-a*^+^* and prc-a^-^ mice.

The third hypothesis suggests that the tension generated by mechanical interactions between fibers running deep through the tissue, such as axons or glia, creates elastic constraints that direct the distribution of the more fluid outer layers (29). Biomechanical studies showing that developing axon tracts are under tension (30) support this hypothesis. The relevance of this hypothesis for cerebellar foliation has been recently demonstrated by an elegant experimental study from the Joyner group (31). Our results, showing that the most prominent morphometric differences correlating with the distinct foliation patterns are in the WM, are consistent with this hypothesis.

Endocannabinoid signaling and CB1 receptors play a role in axon growth and fasciculation (32–35). CB1 is highly expressed in the shafts of developing axons (32) rather than predominantly in the growth cones like the classical axon guidance receptors (36). In axon shafts, CB1 localizes to the periodic actin rings(37), and CB1 activation regulates actin dynamics and axon growth (38–40). It has been proposed that direct adhesive interactions between adjacent axon shafts (zippering-unzippering) may play a key role in the regulation of axon fasciculation-defasciculation (41). Future studies should investigate whether developmental downregulation of CB1 in the axons of the cerebellar contributes to their unzippering, defasciculation, and, consequently, branching of the developing axon tracts, allowing innervation of distinct targets and leading to the formation of secondary lobules.

### Implications and Limitations

Anatomical sex differences in the cerebellum - not a classically recognized sexually dimorphic brain region – suggest that foliation patterns may be influenced by genetic sex, possibly contributing to sex-related anatomical differences in multiple brain regions. Neurosurgeons and radiologists use the patterns of cortical folding as anatomical landmarks; therefore, a better understanding of the variability between individuals in the prevalence of specific fissures and the differences in that prevalence between sexes is beneficial for clinical applications. CNN model reports highly predictable anatomical sex differences in humans but is agnostic to the specific anatomical features that contribute to these differences. Using backpropagation to highlight the pixels most significantly contributing to the categorization could be a viable approach to address this limitation in the future. Anatomical analysis of the mouse cerebella is amendable to a much finer visualization of small anatomical details than human MRI scans, promoting the identification of a correlation between the presence of secondary fissures and the differences in WM shapes. Whether differences in axon fasciculation and branching in the developing WM directly contribute to the differences in the folding of the cortical surface could be addressed by future studies. Utilizing a cross-species approach allowed us to bridge multiple levels of analysis of the mechanisms shaping brain development, from behavioral to anatomical to molecular. The results reported here highlight the advantages and limitations of using this approach to the study of the age-old fundamental question of the relationship between shape and function.

Notably, cannabinoid signaling is revealed to play a role in the regulation of WM tracts development and cortical folding.

## Materials and Methods

### Mouse Colony

All mice used in this study were maintained on an outbred CD1 genetic background. The CD1 IGS stock was acquired from Charles River, USA (https://www.criver.com/products-services/find-model/cd-1r-igs-mouse?region=3611) and replenished by purchasing additional CD1 breeders from Charles River once a year. A breeding colony was maintained in the vivarium at Indiana University Bloomington with a 12-hour light/dark cycle under conditions stipulated by the Institutional Animal Care and Use Committee. CB1 KO (*cb1* -/-) generated by (42) were acquired from Charles River France. CB1-KOs were kept as an outbreed colony on a CD1 background by breeding heterozygous CB1+/- with CD-1 WTs. First-degree crosses (siblings, parents, etc.) were always avoided.

### Tissue Sectioning and Nissl Staining

Mouse brain tissue was fixed by perfusion followed by overnight post-fixation at 4^0^C in 4% paraformaldehyde (PFA). 20 μm sagittal sections were serially collected using a cryostat (Leica). Nissl staining was performed according to a standard protocol. In short, after PBS and dH2O washes, sections were incubated in 0.1% Cresyl Violet solution, followed by dehydration, clearing in xylene, and mounting in Permount. Images of whole cerebellar sections were obtained using 4x objective and mosaics with an Applied Precision DeltaVision Nikon microscope and Motic EasyScan at IUB Imaging Facility. Figures were prepared in Adobe Photoshop and Adobe Illustrator.

### Immunohistochemistry

For immunohistochemistry brain sections were washed 3 times in 1X PBS and incubated in BSA blocking buffer (5% BSA; 0.5% Triton x100; 1X PBS). Anti-calbindin (Rabbit anti Calb, Swant, #CB38, RRID: AB_2721225) primary antibody was applied overnight at 4^0^C in BSA blocking buffer at 1:3000 dilution. Subsequently, slides were washed 3 times in 1X PBS and secondary antibodies (Alexa Flour 488 from Jackson ImmunoResearch) were applied at 4°C overnight (1:600 in BSA blocking buffer). DraQ5 (Cell Signaling) was used (at 1:5000 in PBS) to visualize cell nuclei. Slides were coverslipped using Fluoromount G (SouthernBiotech).

### Morphological Analysis

Areas of interest were traced from midsagittal sections using free-hand tool in FiJi (https://imagej.nih.gov/nih-image/manual/tools.html). <u>Statistical</u> <u>Analysis</u> was performed in GraphPad Prism 9 (www.graphpad.com). Data was collected in total from 114 two-month-old mice of both sexes from 28 distinct liters. The statistical hypothesis tested the magnitude of differences in the area means between males and females with and without *prc-a*. P-values and 95% confidence intervals of difference were evaluated by 2-way AVOVA, or multiple T tests when appropriate. Specific details of statistical analysis for the different comparisons are included in the figures and the results.

### Mouse Behavior

#### Seed Opening

Two-month-old mice were food-deprived overnight (12 hours) for two days. On the second day, during the daytime, a few sunflower seeds were placed in their home cages to facilitate familiarity with this new food. In the morning after the overnight fasting, the mice were placed in the testing cage with 4 seeds – one in each corner. For each seed, time was recorded from first contact with the seed and until the mouse stopped interacting with the seed. Seeds were weighed before and after the experiment to estimate the quantity consumed. Only the trials in which the seed was opened and at least 75% consumed were included in the analysis. Data for each mouse is the average time to open 2-5 seeds.

#### Statistical Analysis

Statistical analysis was performed in GraphPad Prism 9 (www.graphpad.com). Data was collected from 22 female and 17 male animals (7 liters). The statistical hypothesis tested the magnitude of differences in seed opening latencies between sexes depending on the expression of secondary fissures. P-values and 95% confidence intervals of difference were evaluated by 2-way AVOVA, or multiple T tests when appropriate.

#### Rotarod

Two-month-old mice were placed on an accelerating rotarod (UGO Basile) which accelerated from 4 to 40 rpm over a 5 minutes period. The latency until the mouse fell off was recorded. Mice were given three trials per day with a maximum time of 300 seconds (5 minutes) per trial and were given 15 minutes to recover between trials.

#### Statistical Analysis

The data was collected from 22 female and 17 male mice (7 liters) over a time course of 12 trials (four days). Statistical hypothesis of equivalent latency to fall between genotypes was evaluated by Tukey multiple comparison, and by comparing areas under the curve for all trials.

### Human Behavior

Data was taken from the Human Connectome Project (HCP) database (https://www.humanconnectome.org).

#### Peg sorting

the time required to accurately place and remove nine plastic pegs into a plastic pegboard, including one practice and one timed trial with each hand. Raw scores are recorded as the time in seconds that it takes the participant to complete the task with the dominant hand.

#### Statistical Analysis

Statistical analysis was performed in GraphPad Prism 9 (www.graphpad.com). The statistical hypothesis tested the magnitude of mean differences in performance between sexes, T-test analysis was used to compare the two groups.

### Deep learning

We used midline sagittal cerebellar MRI scans from the Human Connectome Project (HCP, https://www.humanconnectome.org/) (Fig. 2). The convolutional neural network (CNN) architecture was built in Keras. Our dataset consists of 1114 images in total, which were split into 85%,10%, and 5% for training, validation, and testing, respectively. Images were taken of a midline slice of the cerebellum after registering labels from the SUIT atlas to the subject’s space and passed through an intensity normalization process that scaled the images for training (14–16). All pixels outside the label set were zeroed so no confounding information from surrounding tissue would contribute to the prediction. The final image sizes after this process were (1167, 875, 3) pixels (image height, image width, and the number of channels, respectively).

Since most of the data outside the label set was set to zero and did not contribute to the prediction while unnecessarily increasing model size, we cropped the images around the cerebellum, reducing image sizes to (420, 280, 3). To further reduce the model size, we used OpenCV library to make the MRI scans grayscale (i.e., single channel). Our testing shows that this adjustment does not change the performance of the model. Images were passed through a series of 6 2D convolutional filters and combined with pooling layers (filter sizes = 4, 8, 16, 32, 64, 128; kernel size = (3, 3); RELU activation). A dense connection layer with a RELU activation function (50 nodes) followed by the final two neurons making classification using softmax function. The model weights were initialized using Xavier glorot, for faster convergence, the dense layer connection between the 50 and final two neurons was supplemented with a 50% dropout for regularization. The model was optimized by making use of the sparse categorical cross-entropy loss by using the Adam optimizer and a learning rate of 0.001. The model was run for 30 epochs with a batch size of 32 for training. The prediction accuracy was used as the metric for scoring model performance. We make use of checkpoints to store the model with the highest validation accuracy to measure performance on test data.

## Acknowledgments

We are grateful to Jim Powers and the IUB Light Microscopy Imaging Facility, and our wonderful undergraduate research assistants Alexandria Bell, Alexander Buchanan, Wesley Cha, Luis Dominguez, Jacob LaMar, Athanasios Liodos, Thomas Metcalf, and Jonah Wirt.

## Author contributions

Conceptualization: AK, FP, KB, BMP

Methodology: AK, FP, BMP, OP

Investigation: KB, OP, BMP, JH, TM, SB, AE

Visualization: AK, KB, JH, BMP, OP

Supervision: AK, FP

Writing—original draft: AK

Writing—review & editing: AK, KM

## Data and materials availability

All data are available in the main text or the supplementary materials.

